# Causal Interactions between Phase- and Amplitude-Coupling in Cortical Networks

**DOI:** 10.1101/2024.03.19.585825

**Authors:** Edgar E. Galindo-Leon, Guido Nolte, Florian Pieper, Gerhard Engler, Andreas K. Engel

## Abstract

Phase coherence and amplitude correlations across brain regions are two main mechanisms of connectivity that govern brain dynamics at multiple scales. However, despite the increasing evidence that associates these mechanisms with brain functions and cognitive processes, the relationship between these different coupling modes is not well understood. Here, we study the causal relation between both types of functional coupling across multiple cortical areas. While most of the studies adopt a definition based on pairs of electrodes or regions of interest, we here employ a multichannel approach that provides us with a time-resolved definition of phase and amplitude coupling parameters. Using data recorded with a multichannel µECoG array from the ferret brain, we found that the transmission of information between both modes can be unidirectional or bidirectional, depending on the frequency band of the underlying signal. These results were reproduced in magnetoencephalography (MEG) data recorded during resting from the human brain. We show that this transmission of information occurs in a model of coupled oscillators and may represent a generic feature of a dynamical system. Together, our findings open the possibility of a general mechanism that may govern multi-scale interactions in brain dynamics.

## Introduction

Functional connectivity (FC) is a measure of the statistical dependencies between the time series recorded from two neuronal populations or brain regions. Evidence supports the idea that FC reflects the communication at different scales in the brain and is highly relevant for brain dynamics. For electrophysiological recordings, a useful approach is to decompose the signals by their phase and amplitude. These representations have revealed the coexistence of two main modes or mechanisms of communication between brain regions. The first one corresponds to a fixed relation between the phases of two brain sources and is measured by the coherence (1, 2). In the second one, the amplitude or envelope of the signals are correlated, suggesting coordinated excitability fluctuations between areas (2–4). Both connectivity modes have been associated with a broad variety of brain disorders (5) and cognitive processes (6, 7) and for many years the studies of FC have been focused on one or the other coupling mode. Whether both types of connectivity are equivalent or perhaps redundant has been addressed recently in studies showing that these coupling modes can differentiate strongly, especially during presence of stimulation (7–9). Furthermore, patterns of both connectivity modes differ across cortical areas and frequency bands. Although linear measures of phase-phase and amplitude-amplitude correlations may be mathematically equivalent for Gaussian distributed signals (10), this is a condition that may be disrupted by external stimuli, motor execution, or change in brain state. Thus, besides the idea that both coupling mechanisms are related to each other, currently we cannot rule out the possibility that both modes might be two representations of a more general mechanism. However, if both mechanisms uncoupled from each other, this would open up the possibility that they may be related in a causal manner.

Here, we explore the possibility that phase- and amplitude-coupling are related through a causal interaction. One difficulty that we faced when addressing this question is that the statistical dependencies that operationalize FC are defined during a certain time window, which ideally is determined by the frequency band of the underlying neural oscillations. This time window is recommended to be sufficiently long to allow for a significance testing. However, if the time window is much longer than the causal interaction there is a high risk that any causal effects may be blurred. To avoid this issue, we adopted an alternative strategy that consists of defining a parameter of similarity across signals in a multichannel manner. This approach allows us to obtain time-series that reflect how consistent phase and amplitude are across channels. These two time-series, that we called phase-consistency (PC) and amplitude consistency (AC), are defined for each time-sample, on which causality analysis can be applied without restrictions or further assumptions. We applied this approach to data recorded from the ferret brain using custom-made µECoG arrays with electrodes covering a large part of the left cerebral hemisphere. Furthermore, we tested whether the results can be confirmed in MEG recordings of resting activity from the human brain. Finally, we explored in a computational neural-mass-model whether such causality effects can be generated from a common neural mechanism.

## Results

### Time-resolved spatial phase and envelope consistencies

We hypothesized that if a causal interaction between phase and amplitude connectivity modes exists, then this should be in the time scales defined by the signal’s frequency. In this case, time scales of the order of one oscillatory cycle are clearly too short for the standard measures of connectivity (11). To measure a causal relation with fine temporal resolution we defined magnitudes that describe the similarity in phase and amplitude across multiple channels at each time point *t’*. For this purpose, we replaced the standard pairwise window-based approach of connectivity (1, 3, 4, 12) with a multichannel measure of similarity across channels for each time *t’*. For phase, similarity is measured by the phase consistency (*PC*), defined in equation (1) (see *Methods*), which is a function of the instant phase *ф*_*k*_(*t*) of channel *k* at time *t* and *N,* the total number of channels. Analogously, for the envelope we used the inverse of the coefficient of variation, defined in equation (2) (see *Methods*), where *μ*(*t*) is the envelope’s mean and *σ*(*t*) is the standard deviation. Figure 1B displays example recording traces of 9 channels located over the visual cortex and filtered at 1-2 Hz (top) and their corresponding *PC* and *AC* time series (middle and bottom rows, respectively).

**Figure 1.**
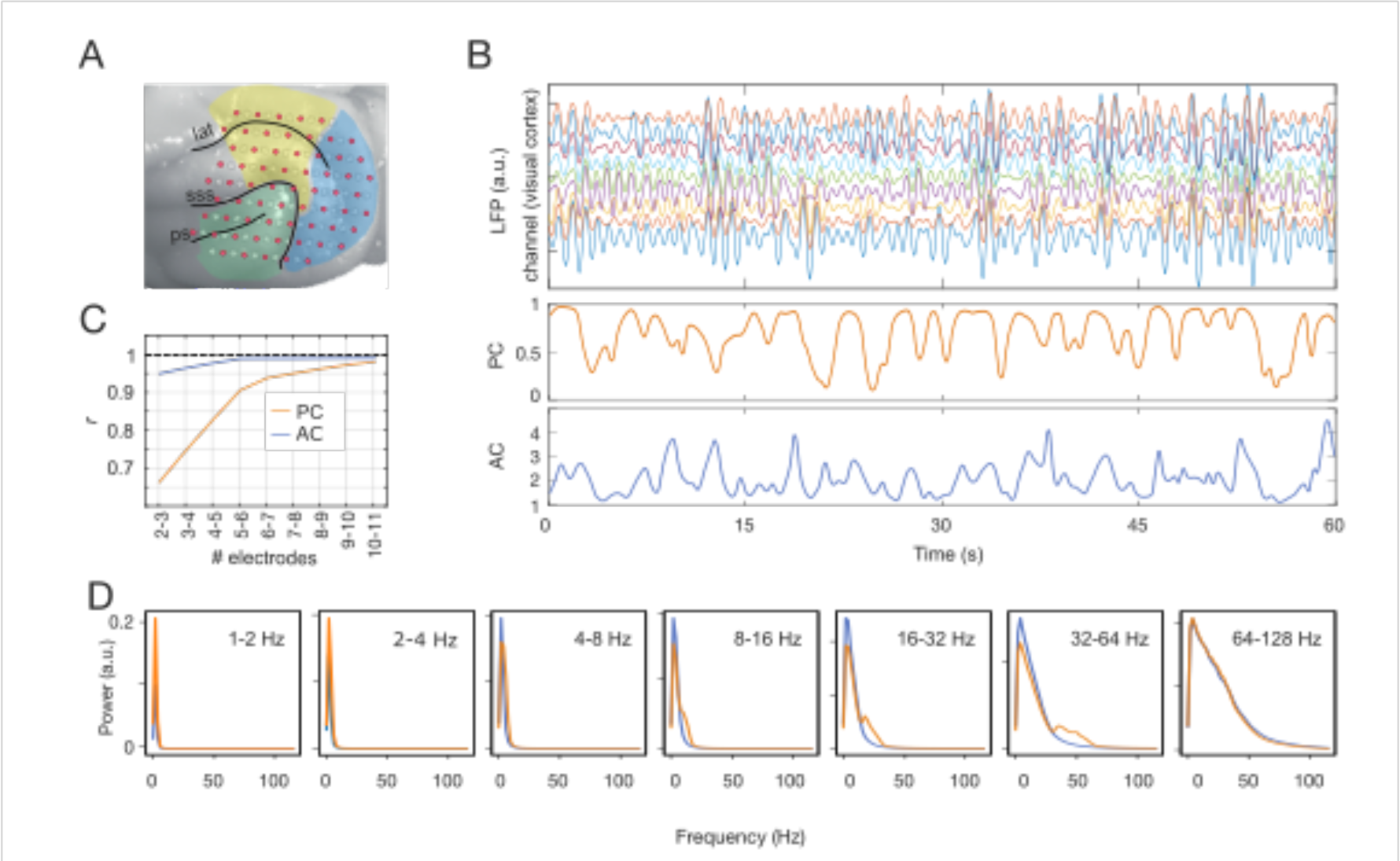
Quantification of PC and AC in ferret LFP data. (A) Representation of our custom-made µECoG array on the left hemisphere of the ferret cortex. Colors indicate the three functional systems: visual (blue), auditory (green), parietal (yellow). (B) Top: Traces of LFP signals (2-4 Hz) of 9 channels located over the visual cortex. Middle: Phase consistency (PC) associated to the above signals. Bottom: associated amplitude’s consistency (AC). (C) The number of electrodes in PC and AC was chosen based on the similarity (correlation r) between the distributions with N and N+1 electrodes. Between 9 and 10 electrodes the similarity was significantly strong. (D) Spectral characteristics of PC (orange) and AC (blue) for different frequency bands of the underlying oscillatory signals.

Clearly, time series, as defined in equations (1) and (2), are sensitive to the size *N*. In our study the selection of *N* is a trade-off between stability in the probability distributions of *PC* and *AC* and the number of channels. One of our goals was to perform the current analysis of the ferret LFP data for two distributions of channels located either within the same functional cortical system or in different systems. Therefore, we selected the smallest number of channels *N* for which the difference of the probability distribution was small compared with the distribution obtained for a *N*+1 population. In this manner we could establish the minimum number of channels required for the within-system condition. Figure 1C shows that the addition of one channel in the population becomes less relevant as the population increases, and that the correlation between *N*=9 and *N*=10 is highly significant (*r*>0.97). Hence, further analyses on the ferret data were performed taking *N*=9 electrodes.

Finally, we asked whether *PC* and *AC* conserve the spectral characteristics of the preprocessed signal. We found for all frequency bands that in general PC and AC spectral distributions were at lower frequencies than the frequency band from which they are originated (Fig. 1D), implying that PC and AC act as low-pass filter. Any component in PC or AC with higher frequency than the LFP signal would suggest the presence of artifactual events.

### Joint probability distribution of PC and AC

To characterize the statistical dependencies between both quantities we calculated the joint probability distribution *p*(*PC*; *AC*) across frequency bands for the ferret LFP data. First, we considered electrodes placed over visual areas according to the anatomical maps ((13) and Fig. 1A) and divided the corresponding *PC* and *AC* time series in intervals of 10 seconds. The resulting probability distribution across intervals and animals is shown on the top row of Figure 2. The distributions show a clear relationship between measures that is highlighted by the cyan lines, which describe the path along which *AC* is maximal given a certain *PC*. This relation was stronger at low frequencies; however, it was maintained up to 128 Hz. As a control we computed the distributions when intervals were shuffled (Fig. 2, bottom row). Under this condition the time relation within each measure was conserved for each measure, but the time relation between intervals was eliminated. The distribution of AC and PC in the shuffled condition was close to random for all frequencies, supporting the hypothesis of a relationship between both measures. Note that this relationship does not imply causality yet, but describes a mere statistical dependency between both measures.

**Figure 2.**
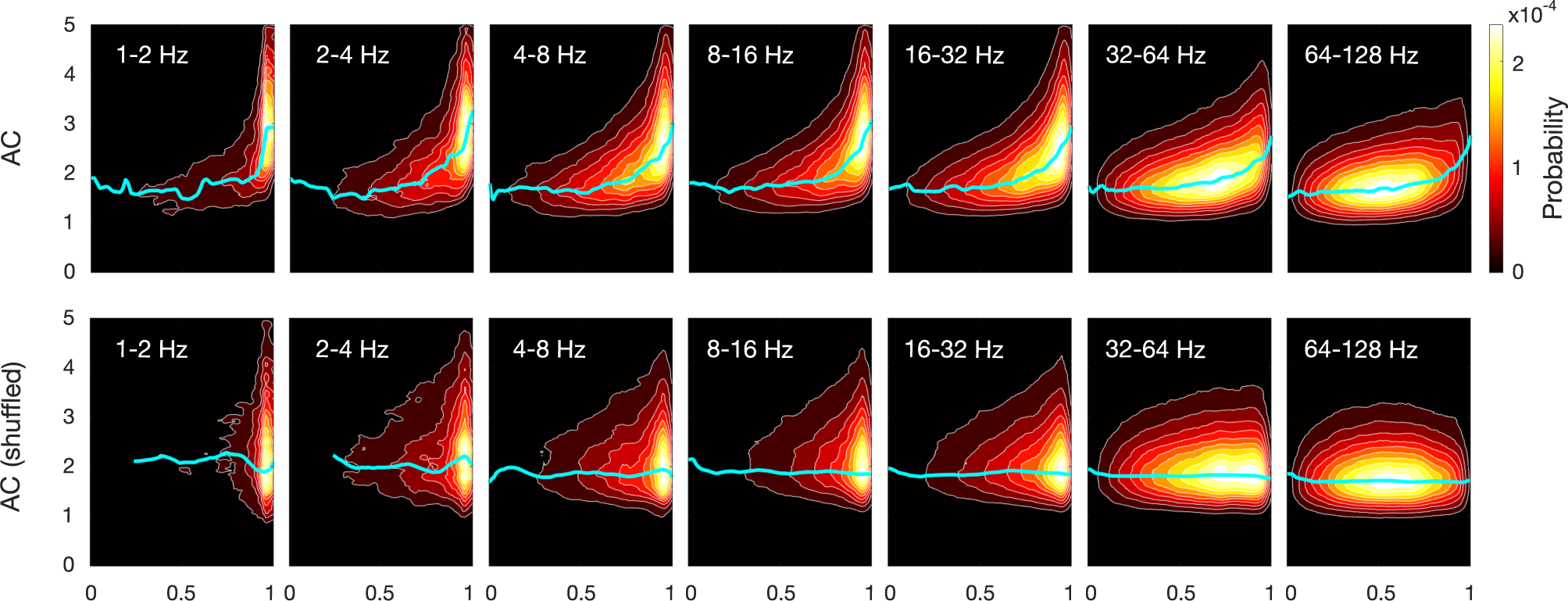
Joint probability distribution of the time series PC and AC in ferret LFP data. To build these distributions we took 100 time-windows of 30 seconds each for all animals. Cyan lines describe the highest probability of AC given a PC. The distributions in the top row suggest a relation between PC and AC that extends to all frequency bands. Bottom row shows the distribution after shuffling the time-windows of both time-series. Note that the cyan line is almost flat, showing the absence of a relation.

### Causal relation between phase- and amplitude consistencies

We hypothesized that the observed correlation might be more than a fortuitous statistical dependency of multichannel phase and amplitudes and, rather, indicate a causal interaction between both coupling modes. To test this hypothesis, we measured causality between *PC* and *AC* using transfer entropy (TE), a model-free method that is valid for non-parametric distributions and that has been used previously in neuroscience to estimate connectivity (14, 15). Again, we extended our analysis across frequency bands, as defined in the previous section. To rule out effects of the electrode selection, we calculated the causality considering electrodes placed over visual areas, over auditory areas and globally distributed across all three functional systems. We contrasted both directions in the causality, i.e., assuming that *PC* is the leader and *AC* is the follower (*PC* → *AC*) and the inverse condition (*AC* → *PC*). The dots in the distributions (Fig. 3) represent the TE during intervals of 30 seconds. Intervals were non-overlapping and randomly selected. We evaluated statistical significance against surrogate data for each interval individually (15–17). The results show a consistent asymmetry in causality dominated by the flow of information *AC* → *PC*. This result was statistically significant in frequency bands below 8 Hz (*p* < 10^-4^, *N* = 343). For higher frequencies (8 to 128 Hz) the flow of information in both directions was not significantly different from the random condition at all.

**Figure 3.**
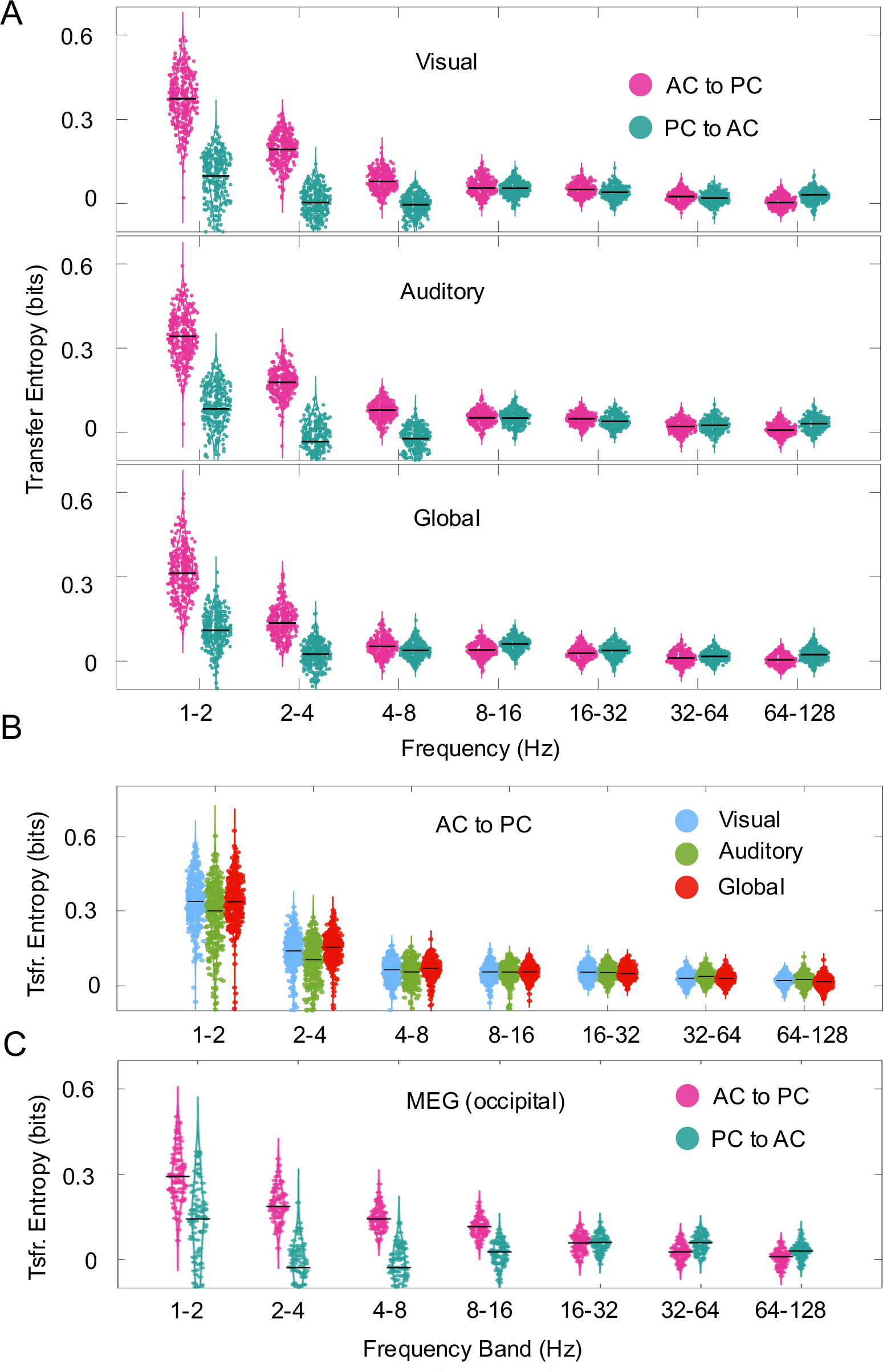
Causal relations *AC* → *PC* and *PC* → *AC*. The causal relation was measured by TE on interval basis (each dot represents an interval of 30 seconds). Horizontal bars show the median TE. (A) TE for different frequency bands in ferret LFP data. Top and middle rows represent conditions where we considered electrodes only over the visual and auditory cortex, respectively. Bottom row represents the sparse condition where we took electrodes from areas regardless of the functional system. We observe a prominent leader role of amplitude coupling (cyan color) with respect to the opposite condition, in particular at low frequency bands (<8 Hz). (B) *AC* → *PC* side-by-side comparison of between the three functional distributions for the ferret data. (C) Transfer entropy for *AC* → *PC* and *PC* → *AC* in human MEG during resting state. AC and PC were obtained from 9 sensors at the occipital region of the left hemisphere.

In the same analysis we asked whether the causality relations are specific to the electrode distribution or the cortical system. Similar patterns of distribution were observed if electrodes were located only in visual areas, only in auditory areas and globally distributed (Fig. 3B). Altogether, our results show a similar causal relation between global network properties based on phase and amplitude coupling modes, suggesting that the mechanisms involved are inherent to the dynamics of the neural networks and not specific to a particular functional brain system.

The above observations might be related to the recording approach, the spatial scale of the signals and the species under consideration. We therefore explored the validity of our results in magnetoencephalography (MEG) data recorded in humans during the resting state (Fig. 3C). We selected 9 sensors located above the occipital cortex of 10 healthy subjects. The selection of the frequencies of the filters were defined as for the ferret LFP data and the duration of the interval for TE computation (dots in Fig. 3C) was also 30 seconds. Interestingly, despite the strong differences in nature and spatial scale of the signals, the asymmetry of TE for AC to PC vs. PC to AC observed in the MEG recordings resembled closely the results obtained for the ferret data. However, this asymmetry extended up to 16 Hz, in contrast to 8 Hz of the ECoG signal. Thus, our results may reveal a fundamental mechanism that is independent of the spatial scale, the recording approach or the species.

### Causal relation is brain-state independent

As we showed in a previous study (18), the short duration of the sleep-wake cycle in the ferret allows the recording of several brain-states within a single recording session. We asked whether the different types of causality observed above were associated with functional states of the brain. For this we defined the asymmetry ratio 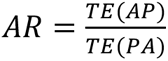 for all intervals by which at least one of the causalities was significant. Thus, a ratio *AR* > 1 means that an alignment in amplitude across channels leads to an alignment in the respective phases, whereas *AR* < 1 means the opposite direction. A value *AR* close to 1 means bidirectionality. In addition, we classified the recordings in three main states using their spectral properties, as detailed in *Methods*. In general, statistical analysis showed no significant evidence that directionality in the causality might be related to the putative brain state (Kruskal-Wallis H test; p>> 0.1). This result extended to all frequency bands and distributions of electrodes in the cortex.

### Information transfer delay

Once demonstrated that the spatial consistency in amplitude may influence the spatial consistency in phase, we proceeded to determine the timescale for this interaction. In its original notation (14), transfer entropy between two random processes is defined as *TE*(*X* → *Y*) = *I*(*Y*_t+1*_; *X*_t_|*Y*_t_), where *t*=1 is the delay between leader and follower. The true interaction delay between variables X and Y is equivalent to the reduction of uncertainty in Y when considering the past values of both Y and X, compared to considering the past values of Y alone (19)

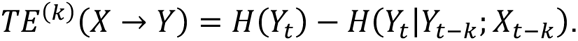

We let the parameter *k* vary between 1ms and 5 seconds and determined the interaction delay *δ* between both time series as

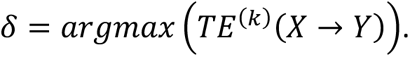

This analysis was separately done for the 3 frequency bands between 1 and 8 Hz (Fig. 4). First, we found similar curves that describe the transfer entropy as function of the delay, with maxima at 208 ms (1-2 Hz), 180 ms (2-4 Hz) and 176 ms (4-8 Hz). In all three bands, the delays correspond to half to one cycle of oscillation, suggesting a mechanism that occurs within a single oscillation cycle and sets an upper limit in the time-scale of the interaction.

**Figure 4.**
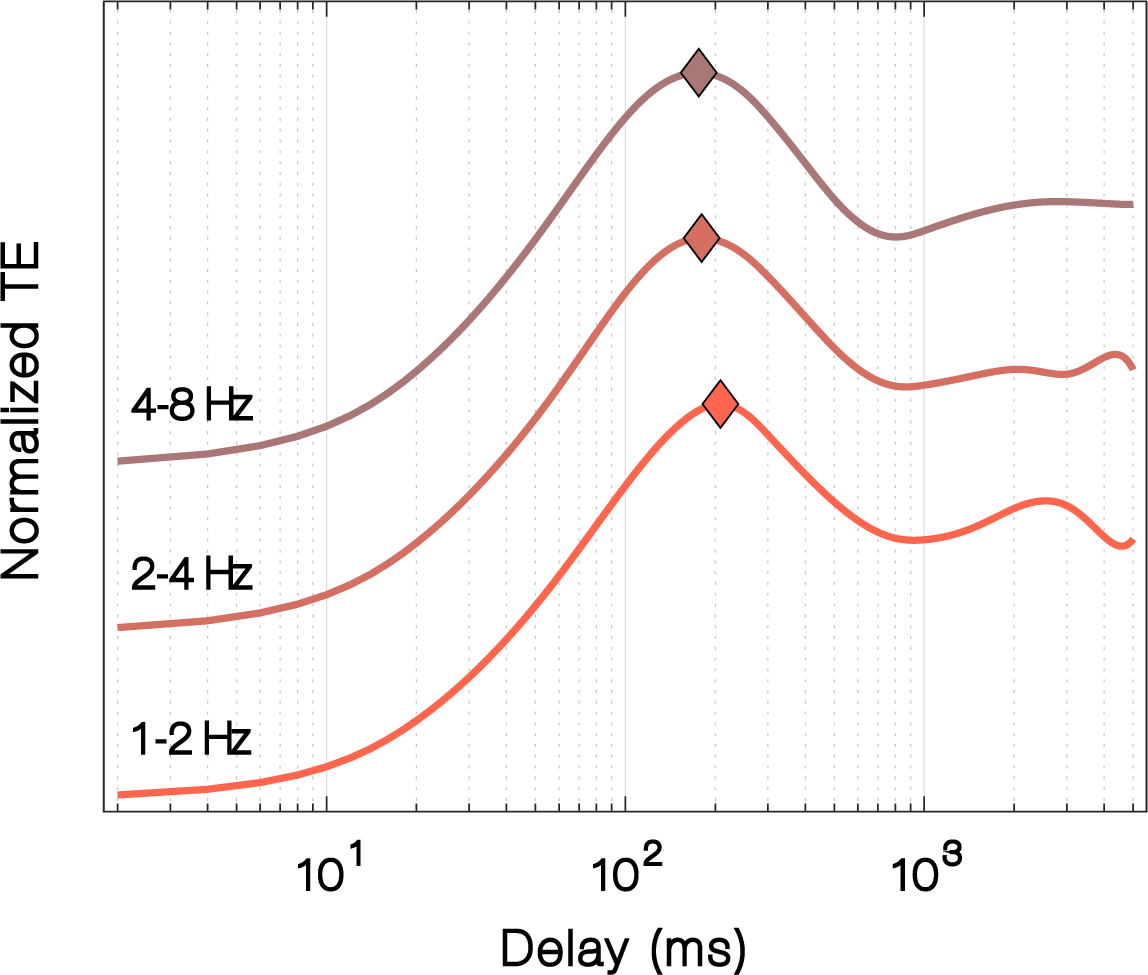
Time relation of the AC to PC causality. The figure shows the normalized transfer entropy between AC and delayed PC in different frequency bands for ferret LFP data.

### Computational simulation in a neural mass model

Finally, we explored whether the characteristics of multichannel coupling and information transfer, as shown above, are exclusively associated to biological processes or rather may reflect a more fundamental property of a dynamical system. To address this question, we used a model based on a non-biological approach. Our rationale here was, if a non-biological system is not able to reproduce characteristics such as the joint distribution of *PC* or *AC*, or the transfer of information, there is the possibility that the observed phenomena have their roots in the complexity of a biological network. On the other hand, if such characteristics were reproduced by a non-biological model, then the transfer of information might represent a general network property.

We used a simple model which consisted of an array of 9 masses coupled by Hooke’s law. We allowed interactions between all couples of masses, mimicking cortical connectivity in a highly simplified manner. The model does not explicitly include parameters such as synaptic delays or excitation and inhibition. However, one could interpret the amplitude of the oscillation of each mass as representing a local field potential. Example traces of simulated signals and the corresponding *PC* and *AC* are shown in Figure 5A. Our model predicted a joint probability distribution of *PC* and *AC* (Fig. 5B) that resembled the experimental data (Fig. 2) with maximal probability at *PC_max_* = 0.96 and *AC_max_* = 3.7. Interestingly, this simple model not only reproduced the causal effect but also the direction of the causality, assigning the *AC* the role of the leader and *PC* the role of the follower as in the low frequency bands in neurophysiological data.

**Figure 5.**
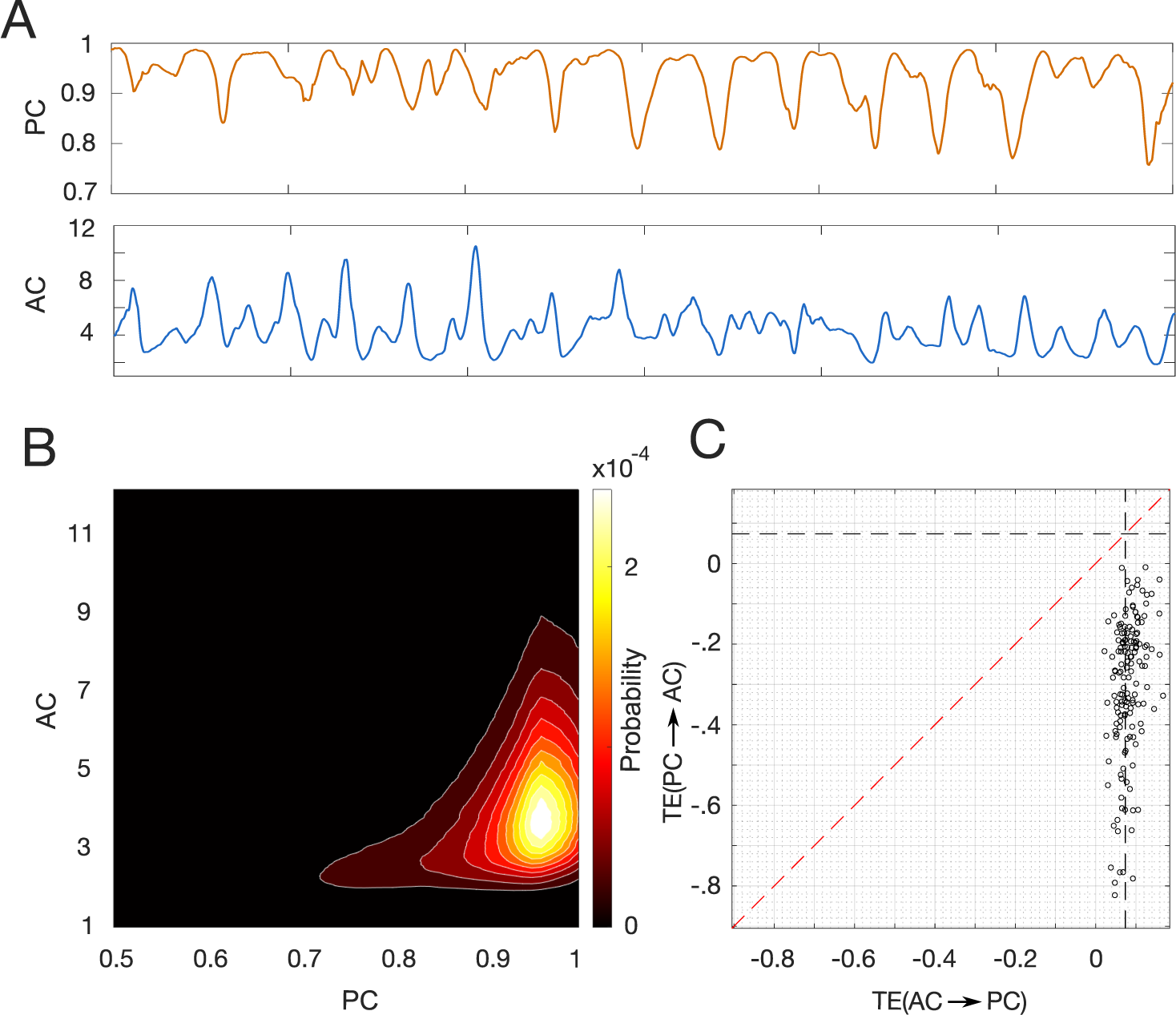
Computational model. For the simulation we considered a set of 9 coupled oscillators allowing pairwise all-to-all interaction. (A) Traces of *PC* and *AC* obtained for simulated traces of coupled oscillators. Note the similarity with the experimental data (Fig. 1B) (B) Joint probability distribution. The similarity with the experimental result is remarkable. (C) Contrast of TE between both directionalities. Our simulation reproduced the leading role of AC to PC.

## Discussion

We have demonstrated that the two major modes of FC that are known to exist in cortical networks, namely, phase- and amplitude-coupling of neural signals (2, 6), interact with each other in a causal manner in ongoing brain activity. Our study was carried out in awake freely moving ferrets using a custom-made µECoG array (7, 18) that allows distributed recordings over large cortical networks with small-sized electrodes that capture rather localized activity, presumably of only few cortical columns (20). In addition, we analyzed the relation between phase- and amplitude-coupling in human resting state data recorded with whole-head MEG (4, 8).

Modes of phase and amplitude interaction have been studied for many years mostly in the context of cross-frequency coupling. In its most common version, the modulation of the amplitude at high frequencies by the phase of low frequency components is quantified (21). This type of interaction, however, describes an instant cross-frequency correlation between different features of signals from the same recording site.

Our study aimed to find a causal relation between different intrinsic coupling modes (2) occurring across distributed areas in the cortex. We introduced gross measures of spatial similarity in phase and amplitude across multiple channels to characterize phase and amplitude’s similarity at each time point *t’*, under the assumption that strong functional coupling results in high similarity of signals across recording sites. First, the interpretation of the resultant time series *PC* and *AC* must be taken carefully. Contrary to standard measures of connectivity, where values of 1 and 0 mean strong and no-coupling, respectively, *PC* may have any value in between even if the signals are perfectly coupled. For instance, if the signals have different phase, but the difference between them remains constant, then *PC* would be a number between 0 and 1 that remains constant over time. Similarly, a low *AC* does not necessarily mean low connectivity. Therefore, we assume that in both coupling modes the mean strength relates inversely with the variability of *PC* and *AC* across time, rather than their instantaneous value. In fact, we observed unexpected transient periods of desynchronization represented by sudden decreases of *PC*. This behavior exhibits similarities to the metastable oscillatory modes (MOMs) recently reported in Cabral et al. (22), which arise at sub-gamma frequencies. These modes are influenced by global parameters such as mean oscillation frequency and mean coupling, rather than the connectomic structure itself.

By means of transfer entropy we observed a causal relation between *AC* and *PC*. This interaction occurs predominantly at low frequencies (1-8 Hz), with amplitude coupling assuming the leading role and phase coupling that of the follower. This result suggests that amplitude coupling, which likely reflects coordinated excitability fluctuations, can have a causal impact on more precise phase coupling in neural networks (2, 6). The leading role of amplitude at low frequencies contrasts with local cross-frequency phase-amplitude coupling, in which the phase at low-frequencies modulates the amplitude of high-frequency oscillations (21).

One of the reasons that motivated the multi-channel approach was the definition of *PC* and *AC* at each time point. We found that the transfer of information occurs with a delay that is inversely proportional to the frequency of the signal, a time scale that is much shorter than the ideal time window used for the measures of connectivity. These time scales are much longer than the synaptic delay within areas, which are typically on the order of a few milliseconds (23, 24). Analyzing the delays relative to one oscillation period, we observed that within the 1-2 Hz band, this corresponds to approximately 2π/3 of the period. For the 2-4 Hz band, this equates to approximately π, and within the 4-8 Hz band, this value increases to approximately 2π. These findings lead us to two conclusions. First, they validate our hypothesis regarding causal interactions manifesting within time scales comparable to oscillation periods, emphasizing the necessity for time-resolved measurements. Second, the observed increase in relative delay at higher frequency bands implies the involvement of components at lower frequencies in the *PC* and *AC*, as illustrated in Figure 1.D.

Our result that similar patterns of causality were observed in ongoing activity recorded by MEG as well is not an obvious at all. First, the neural signal recorded by a single sensor in MEG reflects the activity of a much larger population than the recorded by our µECoG array, where the recorded signals are much better localized and arise from few cortical columns under the recording contact (20, 25). Second, the spatial extent covered by the 9 µECoG-electrodes and the 9 MEG-sensors was quite different; the separation between the most distant recording sites in the ferret was ∼1 cm, whereas in the human data it was ∼6 cm. The coverage proportion of the visual cortex was, however, comparable in both cases. This unveils a mechanism that operates across various scales and broad measures. Another result that reveals the scale-free properties of the mechanism is the similarity of the spectral characteristic for both, *PC* and *AC* across carrier frequency bands (Fig 1.D). The character of the *PC* and *AC* as function of frequency resembles the ∼1/f power-law distribution ubiquitous to self-organized criticality (26).

Although the goal of our study was not the detailed computational simulation of the relation between *PC* and *AC*, we explored to what extent a simplistic, non-biological system can predict the observed phenomena. We started with a model of coupled linear oscillator with noise. Contrary to established models like Kuramoto’s model, our goal was not to explicitly describe effects like synchronization (27). Our model, however, predicts statistical distribution of *AC, PC* and their joint probabilities that resemble the ones observed experimentally. We see two possible scenarios that may explain the transient events of desynchronization in *PC* observed in our model. The first one would be a variation in the delay that is different for each connection (28). This scenario, though, would imply jittering in delay at least comparable with the time scale of the frequency band. For instance, jittering in 2-4 Hz band would require jittering of ∼100 ms to introduce desynchronization. Since changes in the delay are not taken explicitly into account by our model, the fact that the simulation predicts such variation suggests to us that they are due to the close similarity in the spectral properties of the oscillators, which in our case are defined by mass *m_i_* and coupling constants *k_ij_*. Finally, one possibility that may explain why our computational model reproduces the probability distributions and the causal relations is the fact that we considered all-to-all weak interactions.

In conclusion, our results show a causal relation between global representations of phase and amplitude consistency across neural populations. We hypothesize that this relation is an inherent mechanism of brain dynamics and its functional role may be the associated with events that require the coordinated action of global phase and amplitude. In a recent study, Galinsky and Frank (29) in which a network of nonlinear oscillatory propagating modes the emergence of collective synchronized spiking activity was possible only when both, phase and amplitude, were taking into account. The present results clearly indicate a causal impact of amplitude coupling on phase coupling, which raises the question of the putative functional relevance of such an interaction. We have previously suggested that amplitude coupling modes, reflecting correlated excitability fluctuations, may serve to gate, or facilitate, faster phase coupling of neural oscillations (2, 30). This is compatible with recent modeling work indicating that scale-free amplitude fluctuations can have an important influence on phase coupling. Based on these results it has been proposed that critical states of brain networks, as reflected by power-law scaling and long-range temporal correlations, are characterized by intermediate levels of oscillatory coupling, which allow for rapid reconfiguration of functional connections defined by phase relations (31). Resolving the origin and the potential functional implications of interactions between amplitude and phase coupling awaits future research, possibly requiring specific interventions that can separately target these coupling modes.

## Materials and Methods

### Electrophysiological recording of ferret data

Data recorded from 7 freely behaving female ferrets (Mustela putorius furo) were used for the present study. A detailed description of housing, implantation of µECoG arrays and recording procedures have been already reported in (18). Briefly, a custom designed µECoG with 64 electrodes array was implanted over the cortex of the left hemisphere (Figure 1). Before the excised piece of bone was place back in place, we took photographs of the distribution of array contacts on the cortex to assign the position of each electrode to one of 16 anatomical areas based on the map generated by (13). For later analysis we assigned the anatomical areas into 3 main groups comprising visual, auditory and parietal areas, respectively. After recovery (∼7 days) the animal was accustomed to a sound attenuated chamber where the experiments were performed. ECoG signals were digitized at 1.4 kHz (0.1 Hz high pass and 357 Hz low pass filters), and sampled simultaneously with a 64 channel AlphaLab SnRTM recording system (Alpha Omega Engineering, Israel). During the recording sessions the animals were able to move freely, and movements were monitored with an accelerometer mounted to the cable-interface attached to the head. Further information on animal’s preparation, housing and array implantation can be found in (18).

### MEG recording of human resting state data

MEG resting state data were acquired using a 275-channel whole-head system (CTF MEG International Services LP, Coquitlam, Canada) in a magnetically shielded chamber. For this study, we used resting-state measurements in 10 healthy adult subjects, each of whom maintained the eyes closed for an approximate duration of 10 minutes. Prior to their involvement, all participants provided their informed consent in written form. The local ethics committee (Ethik-Kommission der Ärztekammer Hamburg) granted approval for all employed methods, and all procedures were executed in strict adherence to the stipulations of the Declaration of Helsinki. The data utilized in this study were originally collected for a distinct research project. Data acquisition was conducted at a sampling rate of 1200 Hz, and data were offline down-sampled to 300 Hz for subsequent analysis.

### Analysis of ferret data and brain state classification

All data analysis procedures were implemented either with MatLab (Mathworks, Natick, MA) or in Python. All intervals in which recorded signals were larger than 10 standard deviations were considered as noise and consequently rejected for further analysis. Each signal was re-referenced by subtracting the average signal across all 64 channels to remove potential artifacts and reduce effects of volume conduction. Signals were filtered in steps of 2 Hz with a butterworth filter (2th and 4th order). To reduce dimensionality for the analysis, the filtered signals were gathered into 7 frequency bands ranging from 1-2, 2-4, 4-8, 8-16, 16-32, 32-64 and 64-128 Hz, respectively. Within each frequency the resultant signal was z-scored to normalize amplitude fluctuations across frequency bands and electrodes.

Ongoing behaviors display fluctuations in power spectral characteristics that often can be associated with changes in states of the brain. To classify brain states we used a procedure that we have applied previously (18). We performed a power spectral analysis for all individual channels on sliding time-windows (30 seconds) and calculated the global mean spectrogram. This procedure was repeated in sliding windows of 5 seconds. Subsequently, a principal component analysis (PCA) was performed to reduce dimensionality and, finally, a clustering analysis was applied (k-means) to extract 3 main clusters that we identified as distinct brain states. The number of states was based on the quality of different number of clusters, as we showed in a previous study (18).

### Analysis of MEG data

In order to make a fair comparison with the ferret data, we selected a cluster of 9 sensors located in the most occipital region of the left hemisphere. The signals were down-sampled to 300 Hz and then filtered into 7 frequency bands as follows: 1-2, 2-4, 4-8, 8-16, 16-32, 32-64, and 64-100 Hz. Within each frequency band, the resultant signal was z-scored to normalize amplitude fluctuations across electrodes. Brain-state analysis was not performed in the human data as the recording duration was too short for this purpose.

### Time resolved phase- and amplitude consistency

Standard measures of connectivity, such as amplitude correlations or coherence, are defined for pairs of recording sites (e.g., electrodes or brain regions of interest) during a time-window which can vary in duration from sub-seconds to minutes. However, for the study of the causal relation between phase and amplitude coupling modes we considered it convenient to introduce instantaneous measures that can be defined at each time *t’*, rather than for an extended time window. Here, we propose an approach based on the assumption that strong phase (amplitude) coupling is reflected in a less variable relation of phases (amplitudes) across sets of recording sites. Conceptually similar to the phase-locking value (PLV), which reflects how consistent the phase related to a particular stimulus event is across trials (12), we took signals of multiple electrodes and measured the consistency of the phase relation across recording sites defined as:

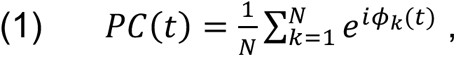

with *ф_k_*(*t*) the phase of channel *k* at time *t*, and *N* the number of channels. This value by itself does not say much about the connectivity. Assuming that the there is a perfect phase correlation between the signals, the *PC* can take any value between 0 and 1. Applying the same rationale to the signal amplitudes, we defined amplitude consistency (*AC*) as:

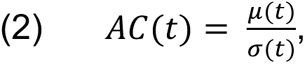

with *μ* t the mean and *σ* the standard deviation at time *t*. Instant phase and amplitude were computed after applying the Hilbert transform to the filtered signals. Note that a consistent phase relation between channels can take any arbitrary value between 0 and 1, and whether the phase-relation remains constant (as an indicator of connectivity) is indicated by a low temporal variation of PC.

### Causality measure

We used transfer entropy (TE) to determine the causal relation between phase-based (*PC*) and amplitude-based (*AC*) time series. TE measures the directed transfer of information between two non-parametric time-series *X* and *Y*, and is defined as the information shared between the past of *X* and the present of *Y* present, given *Y’s* past (14):

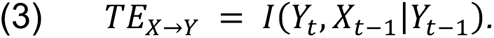

Here we calculated TE in both directions, namely, *PC* playing the role of leader and *AC* the follower, and vice versa. Both quantities were calculated separately for non-overlapping time-windows of 30 seconds and for all frequency bands. The statistical significance test was based on the null-hypothesis of no source-target interaction (16, 32), with time-randomized leader and follower. These surrogates were created from the same set of observations (leader’s interval) maintaining the same distribution but with temporal dependency of the source destroyed. For each individual interval we selected the significance level as the mean plus two times the standard deviation. We selected the highest significance level across intervals and animals. TE was calculated in Python using the open-source implementation PyIF (33).

### Computational simulation

To study whether a relation between *PC* and *AC*, if any, is caused by some biological effect or not, we simulated a non-biological system of coupled oscillators or springs, in which the interaction obeys Hooke’s law:

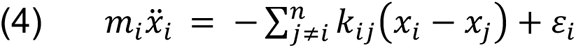

where *m_i_* is the mass, *k_ij_* the coupling constant between nodes *i* and *j*, and *ε_i_* is a noise factor. In the model, we allowed all-to-all interactions between oscillators, rather than only first neighbors. To directly compare with the experimental results, we simulated a system of 9 oscillators randomizing the parameters in each run. Parameter’s estimation *k_i_*, *m_i_* and *ε_i_* was achieved after applying a modest amount of hand-tuning that delivered reasonable predictions. We implemented the following set of parameters: *k_i_* = 3 ± 0.1, *m_i_* = 4 ± 0.2 and *ε_i_* = 0 ± 0.5. We solved the system of coupled differential equations using the function ODE45 in MatLab.

## Acknowledgments

This research was supported by the DFG (SFB936-178316478-A2/Z3, SPP1665-EN533/13-1, SPP2041-EN533/15-1) and by the European Union (project cICMs, ERC-2022-AdG-101097402). Views and opinions expressed in this paper are those of the authors only and do not necessarily reflect those of the European Union or the European Research Council. Neither the European Union nor the granting authority can be held responsible for them.

## Author Contributions

EEGL, FP, and AKE conceived and designed experiments. FP surgically implanted arrays. FP, EEGL and GE performed experiments. EEGL and GN analyzed data. GE wrote the animals ethics permission. EEGL and AKE wrote the paper.

